# Sex differences in the functional network underpinnings of psychotic-like experiences in children

**DOI:** 10.1101/2024.04.22.590660

**Authors:** Elvisha Dhamala, Sidhant Chopra, Leon Q.R. Ooi, Jose M. Rubio, B.T. Thomas Yeo, Anil K. Malhotra, Avram J. Holmes

## Abstract

Psychotic-like experiences (PLEs) include a range of sub-threshold symptoms that resemble aspects of psychosis but do not necessarily indicate the presence of psychiatric illness. These experiences are highly prevalent in youth and are associated with developmental disruptions across social, academic, and emotional domains. While not all youth who report PLEs develop psychosis, many develop other psychiatric illnesses during adolescence and adulthood. As such, PLEs are theorized to represent early markers of poor mental health. Here, we characterized the similarities and differences in the neurobiological underpinnings of childhood PLEs across the sexes using a large sample from the ABCD Study (n=5,260), revealing sex-specific associations between functional networks connectivity and PLEs. We find that although the networks associated with PLEs overlap to some extent across the sexes, there are also crucial differences. In females, PLEs are associated with dispersed cortical and non-cortical connections, whereas in males, they are primarily associated with functional connections within limbic, temporal parietal, somato/motor, and visual networks. These results suggest that early transdiagnostic markers of psychopathology may be distinct across the sexes, further emphasizing the need to consider sex in psychiatric research as well as clinical practice.

## Introduction

Psychiatric illnesses are often preceded by premorbid and/or prodromal states that emerge months to years before the onset of a diagnosable illness[1]. These states include subtle changes in mood, behavior, thought, and cognition that do not meet the threshold for a formal diagnosis. Psychotic disorders manifest across a spectrum ranging from psychotic-like experiences (PLEs) to clinical high-risk states, first episode psychosis, and chronic psychotic disorder[2]. PLEs represent the earliest potential manifestations of psychosis and tend to emerge during childhood and adolescence. PLEs include a range of sub-clinical symptoms that resemble aspects of psychosis, such as perceptual distortions (i.e., hallucinations), false beliefs (i.e., delusions), and disorganized thinking or speech. As many as 60% of youth report PLEs[2, 3], and these experiences are associated with distress and can hinder social, academic, and emotional development[4]. While only a subset of children with PLEs develop psychosis, many develop other forms of psychiatric illnesses, including affective disorders (e.g., depression) and post-traumatic stress disorder[5]. This suggests that PLEs may represent early markers of not only psychosis, but transdiagnostic psychiatric illness more broadly.

Sex and gender differences exist in the development and expression of psychiatric illnesses[6], including psychosis[7]. Women (as per their gender, hereafter referred to as ‘women’) and individuals assigned female sex at birth (hereafter referred to as ‘females’) are more likely to exhibit internalizing problems (e.g., loneliness), while men (as per their gender, hereafter referred to as ‘men’) and individuals assigned male sex at birth (hereafter referred to as ‘males’) are more likely to exhibit externalizing problems (e.g., aggression) [8]. Psychosis tends to have an earlier onset in men and males along with more several clinical features relative to women and females[9]. PLEs also differ across sexes and genders such that delusions are more common in women and females while hallucinations are more common in men and males[10]. Some recent studies have examined the neurobiological underpinnings for these phenomenological differences. Functional magnetic resonance imaging is used to indirectly estimate neuronal activation based on changes in blood oxygenation. Regional functional activation signals can be correlated to assess the functional connectivity (or coupling) between brain regions. Disruptions in functional connectivity within the frontoparietal control and default mode networks have been implicated in a wide range of psychiatric illnesses and mental health traits[11-13]. Critically, these functional networks exhibit sex differences[14, 15] and they are uniquely related to transdiagnostic psychiatric symptom domains (e.g., internalizing and externalizing problems) across the sexes in youth[16]. However, it remains to established whether PLEs map onto shared or distinct functional networks across the sexes. An analysis of the sex-specific functional network underpinnings of PLEs in children will reveal early brain-based markers indicative of potential future onset of psychiatric illness that can be used to develop targeted preventative strategies.

Here, we sought to characterize the sex-specific functional network underpinnings of PLEs in children. To do so, we used multivariate analyses to quantify the functional networks associated with the PLEs in a large sample of children from the Adolescent Brain Cognitive Development (ABCD) Study (n=5620, 2671 females, ages 9-10) in a sex-specific manner. First, using a brain-based predictive modeling approach, we demonstrate that functional connectivity is significantly associated with the total number of PLEs and the resulting distress in females and males. Next, evaluating whether the functional networks underlying PLEs are shared or distinct across the sexes, we find that although there is some overlap, a distinct set of networks are implicated in PLEs in females and males. Finally, characterizing the functional network underpinnings of PLEs, we reveal that, in females, a diffuse set of cortical and non-cortical functional connections are associated with PLEs, whereas in males, PLEs predominantly map onto limbic, temporal parietal, somato/motor, and visual networks.

## Results

### Males report more PLEs and greater distress relative to females

We assessed sex differences in PLEs in children. PLEs were assessed using the Prodromal Questionnaire-Brief Child Version, which is comprised of 21 questions. Total and Distress scores were computed, consistent with prior literature[3, 17, 18]. In our sample, 57% of females and 60% of males reported one or more PLEs (out of 21), with 18% of females and 20% of males reporting five of more PLEs. Across all females, an average Total score of 2.31±3.43 and Distress score of 5.74±10.40 was reported. Across all males, an average Total score of 2.55±3.54 and Distress score of 5.84±10.06 was reported. Although higher Total (Mann Whitney U statistic, U=3.31x10^6^, p_corrected_<0.01) and Distress (U=3.36x10^6^, p_corrected_=0.03) scores were reported in males relative to females, the observed differences in the distribution of the Total and Distress scores were minimal (**Figure 1A**). These results are in line with those previously reported in the ABCD Study[19], indicating that the subsample included here with complete resting-state functional MRI and behavioral data is representative of the larger ABCD cohort. Of note, these data indicate that PLEs are not uncommon in children.

**Figure 1:**
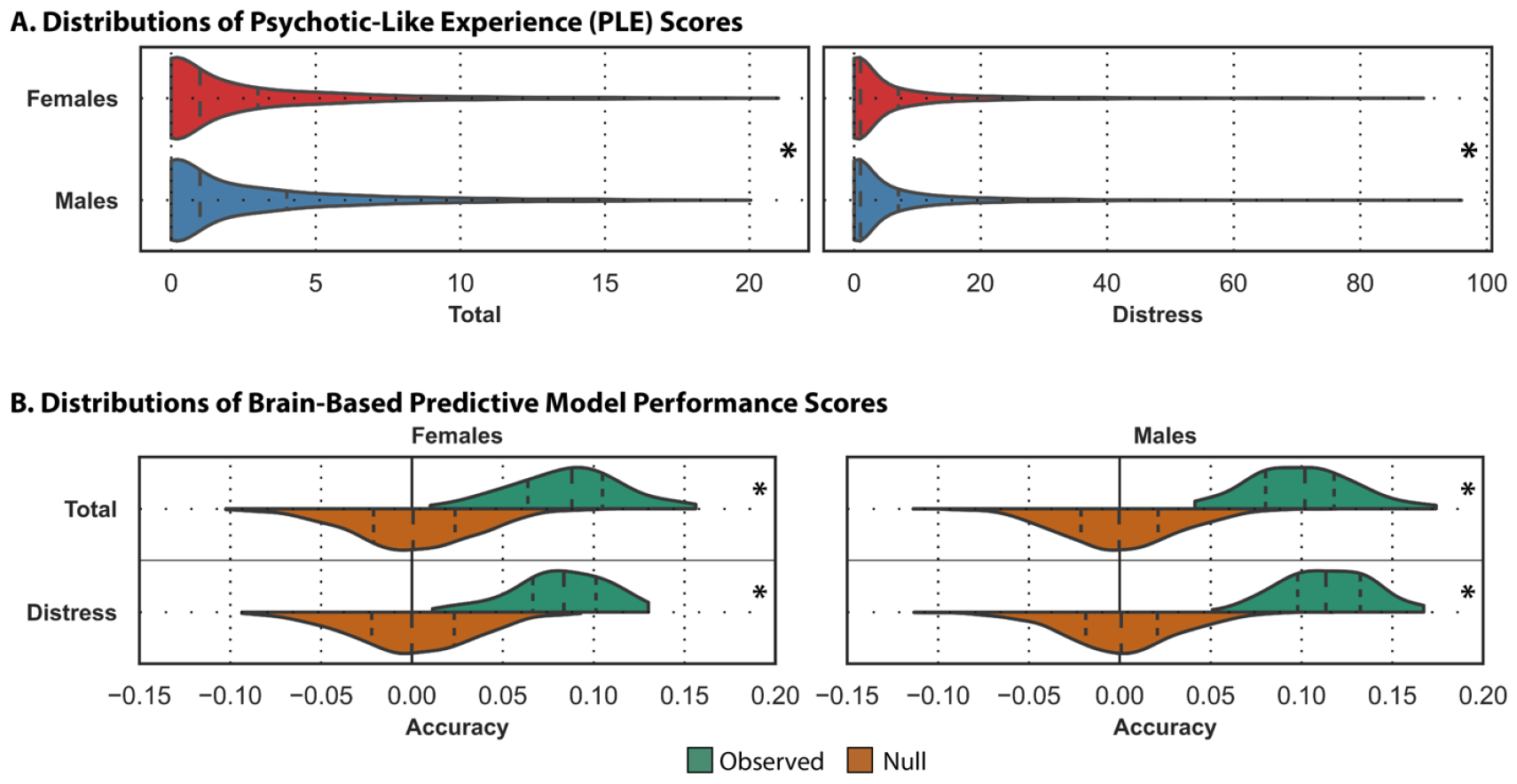
Functional connectivity is associated with PLEs. (A) Violin plots display the distribution of the Total (left) and Distress (right) PLE scores for assigned females (red) and males (blue). Total and Distress scores, although significantly different across the sexes, as denoted by the asterisks (*), exhibited largely overlapping distributions. (B) Prediction accuracy (correlation between observed and predicted values) obtained by the brain-based predictive models trained to predict Total and Distress PLE scores in assigned females (left) and males (right). Predictions for Total and Distress scores in both sexes were significantly better than chance (p_corrected_<0.05), as denoted by the asterisks (*). Results are shown for both the observed (green) and the null (orange) predictions. For all violin plots, the shape indicates the entire distribution of values; the dashed lines indicate the median; and the dotted lines indicate the interquartile range.

### Functional network connectivity is significantly associated with PLEs

Using sex-specific cross-validated linear ridge regression models, we quantified the sex-specific associations between functional network connectivity and the Total and Distress scores for PLEs. As in similar studies using brain-based predictive models to study associations between functional connectivity and behavioral measures in the ABCD sample[13, 16, 20, 21], model performance was evaluated using measures of prediction accuracy (i.e., correlation between observed and predicted scores). Across both sexes, individual variability in functional connectivity was associated with the Total and Distress scores (**Figure 1B**). Although, these observed associations were numerically higher in males for both Total (prediction accuracy, r_Total_= 0.10) and Distress (r_Distress_=0.11) scores compared to females (r_Total_=0.09; r_Distress_=0.08), all brain-based prediction models yielded significant results (p_corrected_<0.05 for all models relative to null models). These results demonstrate that individual variations in functional connectivity can predict both the endorsement of PLEs and the associated distress in youth. As such, functional network connectivity may serve as a potential early biomarker of transdiagnostic psychiatric illness risk.

### Overlapping and distinct functional networks underlie PLEs across the sexes

We applied the Haufe transformation[22] to the model feature weights to increase their interpretability and reliability[20]. As in prior work[15, 16], we averaged the transformed weights across the multiple train/test splits to obtain a mean feature importance score. To assess whether shared or distinct functional connections were associated with PLEs across the sexes, we computed the full correlation between the features extracted from each sex to predict Total and Distress scores (**Figure 2A**). Functional connections associated with Total and Distress scores were strongly correlated within each sex (r_females_=0.97, r_males_=0.97), while functional connections associated with each score were moderately correlated across the sexes (r_total_=0.59, r_distress_= 0.57). Finally, the associations between functional connectivity and the Total and Distress scores were summarized to a network-level by mapping the absolute regional pairwise feature weights onto 17 cortical networks[23] and one non-cortical network. In females, while the strongest associations were observed in temporal parietal, default, limbic, and ventral attention networks, dispersed functional connections throughout the brain were associated with PLEs (**Figure 2B for Distress and S2 for Total scores**). In males, limbic networks exhibited the strongest associations with PLEs followed by weaker associations in temporal parietal, somatomotor, and visual networks (**Figure 2B for Distress and S2 for Total scores**). Summarized network-level associations were also computed (**Figure 3**) and demonstrate similar results. Prior work in this area has identified associations between resting-state network connectivity in control, default mode, and cinguloparietal networks and PLEs[24]. Here, we find that while those networks are implicated to some extent, other networks (e.g., limbic, ventral attention, somatomotor, visual) also contribute to PLEs, and these associations between connectivity and PLEs differ across males and females. Taken together, this suggests that the functional network underpinnings of PLEs overlap to some extent across the sexes in children but also include distinct network contributions.

**Figure 2:**
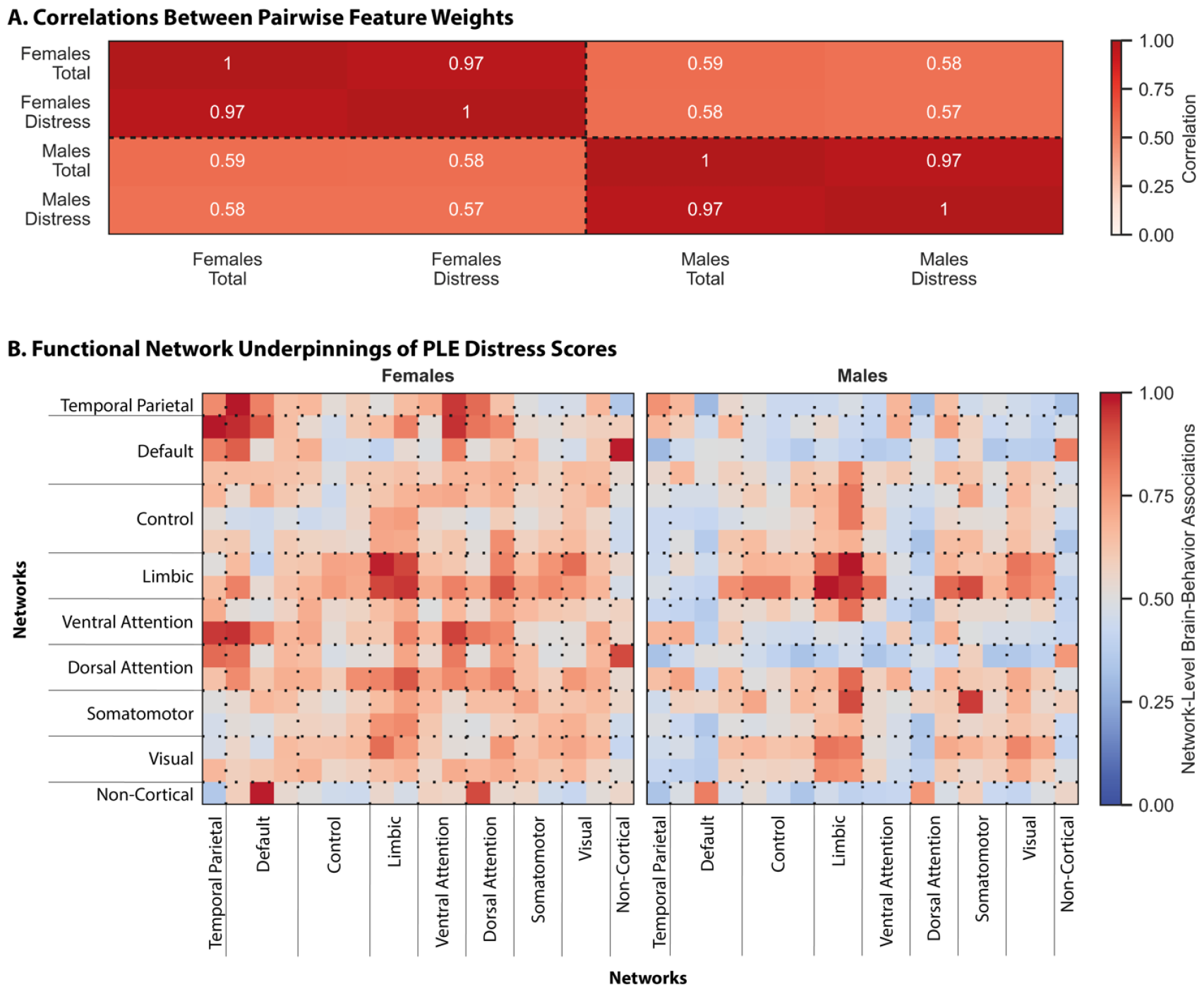
Overlapping and distinct functional networks underlie PLEs across the sexes. (A) Full correlation coefficients between the Haufe-transformed pairwise regional feature weights from models trained to predict Total and Distress PLE scores in assigned females and males. Warmer colors indicate a stronger correlation between the feature weights. (B) Absolute associations between functional network connectivity and PLE Distress scores in assigned females (left) and males (right) are shown as per the colormap. Warmer colors indicate stronger associations and cooler colors indicate weaker associations. To facilitate visualization, values within each matrix were divided by the maximum value within that matrix.

**Figure 3:**
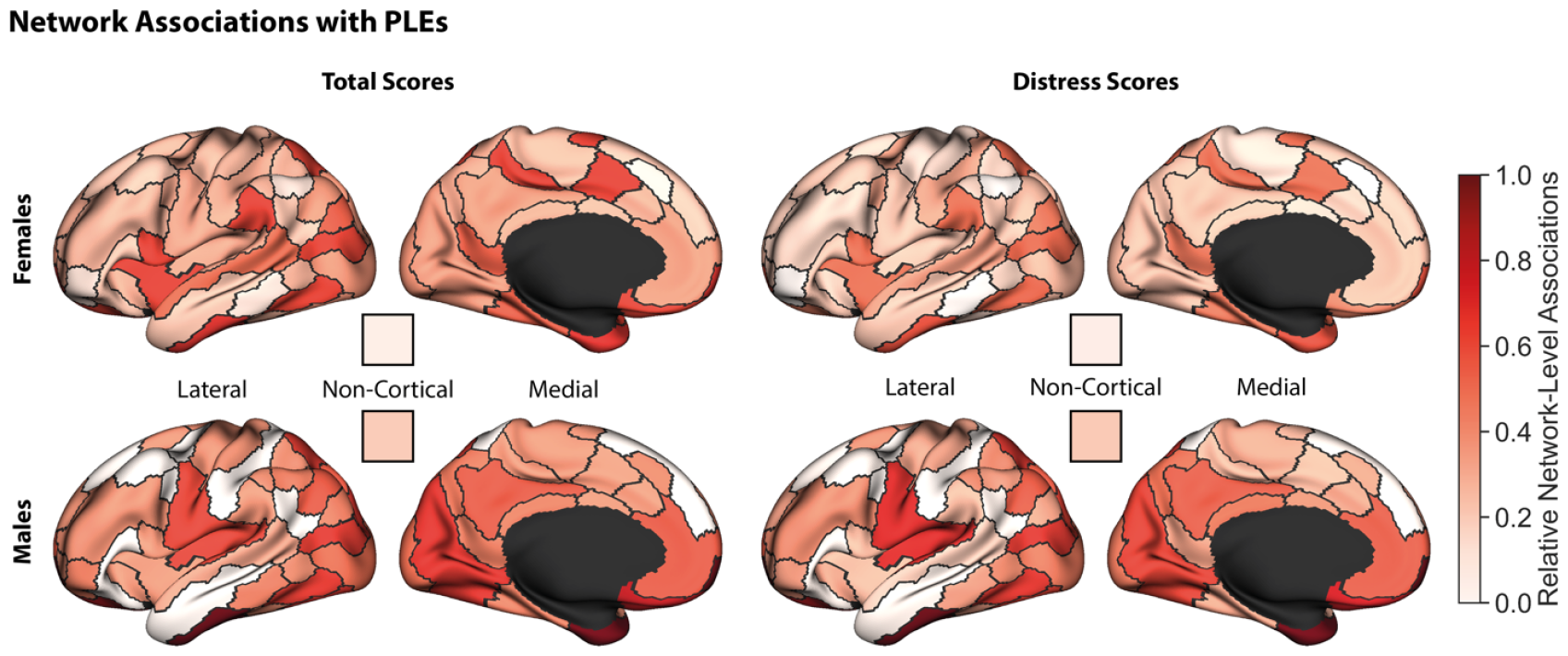
Overlapping and distinct networks are associated with PLEs across the sexes. Summarized relative network-level associations between functional networks and PLE Total (left) and Distress (right) scores in assigned females (top) and males (bottom). Cortical networks are shown along lateral and medial surfaces, and non-cortical networks are visualized as a single square as per the colormap. Warmer colors indicate a stronger relative absolute association between the networks and the scores. To facilitate visualization, values for the network associations were divided by the maximum value for that score across both sexes, then normalized based on the minimum and maximum values for that score across both sexes.

## Discussion

PLEs represent one of the earliest markers of transdiagnostic psychopathology[25]. An understanding of the neural bases of PLEs is critical for the subsequent development of targeted prevention strategies to mitigate the potential negative sequelae of these experiences in at-risk youth. Here, we demonstrate that individual variations in functional network connectivity are significantly and uniquely associated with PLEs across the sexes in children. First, examining the prevalence in PLEs, we confirm prior work, showing that approximately 3 in 5 children report PLEs, and the number of experiences and the resulting distress is largely comparable across females and males. Next, quantifying the relationships between functional connectivity and PLEs, we demonstrate that individual differences in functional connectivity are significantly associated with PLEs in both sexes. Finally, comparing the network underpinnings of PLEs across the sexes, we find that they are moderately correlated but do exhibit notable differences. In females, dispersed functional connections throughout cortical and non-cortical structures are implicated in PLEs, whereas in males, PLEs are primarily associated with functional connections in limbic, temporal parietal, somatomotor, and visual networks. Collectively, our results suggest that the neurobiological underpinnings of PLEs may, in part, be influenced by assigned sex.

Extant literature has shown that PLEs are related to changes in neurobiological structure and function. PLEs have previously been linked to lower cortical gyrification in left fronto-temporal regions and higher cortical gyrification in right parietal cortex[26]. Reduced cortical and subcortical volumes have also been reported in youth with more distressing PLEs[27]. In the functional domain, connectivity between cerebellar and control networks, between cerebellar and cingulo-parietal networks, and within the control network have been implicated in distressing PLEs[27]. A separate study found that connections within the control, default, and cingulo-parietal networks were associated with PLEs[24]. Unfortunately, these studies only considered assigned sex as a covariate, if at all, restricting their ability to capture critical sex differences in the neurobiological underpinnings of PLEs[28]. Our present analyses demonstrate that the overlapping and unique individual variations in functional networks underlie PLEs across the sexes. Specifically, we observe significant associations between functional connectivity and Total and Distress PLE scores in the range of 0.08 and 0.11. PLEs are phenomenologically heterogeneous, so it is not surprising that these effect sizes, while significant, are relatively small in magnitude. Notably, our observed effect sizes are comparable to those reported between gender and risk-taking behavior (r=0.08) as well as combat exposure and PTSD (r=0.11)[29].

Self-reported PLEs are highly prevalent in children and are associated with parent-report psychopathology[3], delayed functional development[27], and future development of psychiatric illnesses[5]. Although not all youth who report PLEs develop psychosis, the risk of conversion is 3.5 to 4 times higher than individuals without PLEs[30, 31]. Moreover, many of them experience other mental health problems, including anxiety, depression, PTSD, substance dependence, and suicide attempts[5]. In fact, a New Zealand birth cohort study suggests more than 90% of children with strong PLEs have some form by clinically diagnosable psychiatric disorder by age 38[5]. This suggests that childhood PLEs may be predictive of poor mental health during adolescence and adulthood. As such, children who experience PLEs may benefit from early interventions directed at improving their mental well-being[32]. Here, we establish the sex-specific neural bases of PLEs in a large sample, providing a critical foundation for subsequent research on the neurobiological mechanisms driving the future emergence of transdiagnostic psychopathology in these individuals. Importantly, the network associations captured here are, for the most part, unique from those previously reported between functional connectivity and other psychiatric illness-linked behaviors (e.g., internalizing and externalizing) in this sample[16]. This suggests the brain-behavior associations captured here are specific to PLEs and do not represent general psychopathology in children. The neurobiological markers of PLEs identified here can subsequently be validated in smaller, disease-specific samples (e.g., NAPLS [33-35]). The present findings can also be used to guide the development of targeted intervention strategies that may modulate these functional connections, ultimately improving resilience.

Sex differences in the neurobiological underpinnings of PLEs, and psychosis more broadly, have been largely understudied to date. In schizophrenia, men and males have a greater risk of developing negative symptoms (e.g., avolition, anhedonia, alogia), while women and females tend to display more affective symptoms (e.g., depression, impulsivity, emotional instability)[9, 36]. Women and females also exhibit stronger cognitive functioning along executive, verbal, and processing domains, while men and males outperform females along memory and attention domains[36]. Women and females are at a particularly elevated risk of developing psychosis during critical hormonal transition periods (e.g., pregnancy, post-partum, menopause)[37], and earlier menarche is associated with a later onset of schizophrenia[38]. Here, we observe critical sex differences in the associations between functional brain networks and PLEs, providing an insight into the neural bases that may, in part, underlie sex differences in psychosis and schizophrenia. The sex differences observed here may be influenced by a number of sex- and gender-related factors, including differences in functional network maturation trajectories across the sexes[39-41]. Additional analyses using the longitudinal ABCD data and other datasets will inform whether the associations observed here remain consistent throughout adolescence and adulthood.

Importantly, the reported findings are subject to several limitations. First, these results were obtained using a single dataset. Although, our sample is relatively large (n=5,260) and we used a cross-validated, sex-specific brain-based predictive modeling approach known to generate reliable results[16], we cannot rule out the possibility that these results may be limited in their generalizability[42]. Future research in global open-access datasets with comparable neuroimaging and behavioral data could analyze whether the reported findings are specific to the ABCD sample considered here. Second, we only considered the influences of binary assigned sex in these analyses. Although sex is not binary, participants in this sample reported their assigned sex as Female or Male, and as such we were only able to compare the network underpinnings of PLEs between the binary sexes. Additionally, gender has been shown to influence functional network organization[15] and the manifestation of psychiatric illness[43].

Unfortunately, gender identity data was not acquired in this sample at the baseline time point and thus was not considered here. It is possible that our results are, in part, driven by gender rather than sex. Subsequent analyzes considering both sex and gender could disentangle their independent and intersecting influences on the neural bases of PLEs. Finally, we assessed the relationships between functional connectivity and PLEs at a single time-point from the baseline acquisition as we sought to identify the earliest neurobiological markers of transdiagnostic psychopathology. However, the networks implicated in these findings mature throughout adolescence[39]. Therefore, it is possible that the brain markers of PLEs, and poor mental health more broadly, shift throughout the transition from childhood to adolescence to adulthood. Future work incorporating multiple time points and a longitudinal analytical framework will be able to assess these potential changes.

Sex is a fundamental part of human identity and can influence both neurobiology and mental health. An understanding of how sex shapes the relationships between neurobiology and mental health is necessary for the advancement of mental health prevention and intervention strategies that are tailored to each sex. Here, we identify unique functional network markers of PLEs across the sexes, revealing sex-specific early markers that may be broadly indicative of poor mental health and forecast the development of transdiagnostic psychopathology.

## Methods

The analyses described here use a similar framework as in our prior work [16, 44-46] to perform novel analyses to establish the functional network underpinnings of psychotic-like experiences in a sex-specific manner.

### Dataset

The Adolescent Brain Cognitive Development (ABCD) dataset is a large community-based sample of children and adolescents who were assessed on a comprehensive set of neuroimaging, behavioral, developmental, and psychiatric batteries[47]. Here, we used minimally preprocessed neuroimaging data and behavioral data acquired at the baseline from the NIMH Data Archive for ABCD Release 2.0.1. Magnetic resonance (MR) images were acquired across 21 sites in the United States of America using harmonized protocols for GE and Siemens scanners. In line with our prior work[13, 48], exclusion criteria were used to ensure quality control. As recommended by the ABCD consortium, we excluded individuals who were scanned using Philips scanners due to incorrect preprocessing (https://github.com/ABCD-STUDY/fMRI-cleanup). For the T1 data, individuals who did not pass recon-all quality control[49] were removed. For the functional connectivity data, functional runs with boundary-based registration (BBR) costs greater than 0.6 were excluded. Further, volumes with framewise displacement (FD)>0.3 mm or voxel-wise differentiated signal variance (DVARS)>50, along with one volume before and two volumes after, were marked as outliers and subsequently censored. Uncensored segments of data containing fewer than five contiguous volumes were also censored[50, 51]. Functional runs with over half of their volumes censored and/or max FD>5mm were removed. Individuals who did not have at least 4 minutes of data were also excluded. Individuals who did not have all of the relevant behavioral data were also excluded. Finally, we excluded siblings to prevent unintended biases due to inherent heritability in neurobiological and/or behavioral measures. Our final sample comprised 5260 children (2571 females, ages 9-10 years).

### Sex

We considered sex in terms of sex assigned at birth (referred to as ‘sex’)[52].

### Psychotic-Like Experiences

We considered the number and severity of psychotic-like experiences as measured by the self-report Prodromal Questionnaire–Brief Child Version. This questionnaire has previously been validated in the ABCD sample [3], and is considered to be a useful measure of early risk for psychosis. Moreover, psychotic-like experiences in the ABCD sample, as measured by this questionnaire, have been shown to be associated with a similar set of familial, cognitive, and emotional factors as in adults with psychosis[3]. Consistent with prior literature[3, 17, 18], we considered Total and Distress scores in these analyses. For each question, participants were first asked if they previously had that experience (1=Yes, 0=No). If they answered yes, they were then asked if the experience bothered them (1=Yes, 0=No). If they again answered yes, they were asked the extent to which the experience had bothered them (1=Not very bothered, 2=Slightly bothered, 3=Moderately bothered, 4=Very much bothered, 5=Extremely bothered). The sum of the endorsed questions corresponding to each PLE (range 0 to 21) was used to compute the Total score, and the total number of endorsed questions was weighted by the level of distress to compute the Distress score (range 0 to 126). Non-parametric Mann-Whitney U rank tests were used to evaluate sex differences in the Total and Distress scores. Resulting p-values were corrected for multiple comparisons across the two scores using the Benjamini-Hochberg False Discovery Rate (q=0.05) procedure[53].

### Image Acquisition and Processing

The minimally preprocessed MRI data were processed as previously described[13, 54]. Briefly, minimally preprocessed T1 data were further processed using FreeSurfer 5.3.0[55-58] to generate cortical surface meshes for each individual, which were then registered to a common spherical coordinate system[57, 58]. Minimally preprocessed fMRI data were further processed with the following steps: (1) removal of initial frames, with the number of frames removed depending on the type of scanner[49] and (2) alignment with the T1 images using boundary-based registration[59] with FsFast. Framewise displacement (FD)[60] and voxel-wise differentiated signal variance (DVARS)[61] were computed using fsl_motion_outliers. Respiratory pseudomotion was filtered out using a bandstop filter (0.31-0.43 Hz) before computing FD[62-64]. A total of 18 nuisance covariates were also regressed out of the fMRI time series: global signal, six motion correction parameters, averaged ventricular signal, averaged white matter signal, and their temporal derivatives. Regression coefficients were estimated from the non-censored volumes. Global signal regression was performed as it has been shown to improve behavioral prediction performance[65, 66]. Finally, the brain scans were interpolated across censored frames using least squares spectral estimation[67], band-pass filtered (0.009 Hz ≤ f ≤ 0.08 Hz), projected onto FreeSurfer fsaverage6 surface space, and smoothed using a 6 mm full-width half maximum kernel. Once processed, we extracted regional time series for 400 cortical[68] and 19 non-cortical[69] parcels. We computed full correlations between those time series yielding a 419x419 pairwise regional functional connectivity matrix. All processing as described was completed on a local server.

### Predictive Modeling

Linear ridge regression models capture robust, reliable, and interpretable associations between neuroimaging and phenotypic data while avoiding data leakage and minimizing overfitting[70, 71]. Here, we used sex-specific cross-validated linear ridge regression models to predict the Total and Distress scores based on functional connectivity in a sex-specific manner. For each sex, we randomly shuffled and split the data into 100 distinct train and test sets (at approximately a 2:1 ratio) without replacement. Imaging site was considered when splitting the data such that we placed all participants from a given site either in the train or test set but not split across the two. Within each train set, we optimized the regularization parameter using three-fold cross-validation while similarly accounting for imaging site as in the initial train-test split. Once optimized, we evaluated models on the corresponding test set. We repeated this process for each of 100 distinct train-test splits to obtain a distribution of prediction accuracy and explained variance. To evaluate model significance, for each set of predictive models, a corresponding set of null models was generated as follows: the output variable was randomly permuted 1000 times, and each permutation was used to train and test a null model using a randomly selected regularization parameter from the set of selected parameters from the original model. Prediction accuracy from each of the null models was then compared to the average accuracy from the corresponding distribution of model accuracies of the original models. The p-value for each model’s significance is defined as the proportion of null models with prediction accuracies or explained variances greater than or equal to those corresponding to the original distributions. All p-values were corrected for multiple comparisons across the two scores using the Benjamini-Hochberg False Discovery Rate (q=0.05) procedure[53].

### Feature Weights

We used the Haufe transformation[22] to transform feature weights obtained from the linear ridge regression models to increase their interpretability and reliability[13, 54, 72]. For each train split, we used feature weights obtained from the model, W, the covariance of the input data (functional connectivity), S_*x*_, and the covariance of the output data (behavioural score), S_*y*_, to compute the Haufe-transformed feature weights, A, as follows: 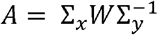. We then averaged the Haufe-transformed feature weights across the 100 splits to obtain a mean feature importance value. We computed full correlations between mean feature importance obtained from the sex-specific models to evaluate whether they relied on shared or unique features to predict PLEs. For all models, we also summarized pairwise regional feature importance at a network-level to support interpretability as previously described[45]. To do so, cortical parcels were assigned to one of 17 networks from the Yeo 17-network parcellation[73], and subcortical, brainstem, and cerebellar parcels were assigned to a single non-cortical network for convenience. Absolute regional pairwise feature weights were averaged to yield network-level estimates of associations between functional connectivity and psychotic-like experiences for each sex. Finally, pairwise network-level were summarized to a network-level by taking the sum across the pairwise network connections.

## Acknowledgements

Data used in the preparation of this article were obtained from the Adolescent Brain Cognitive Development^SM^ (ABCD) Study (https://abcdstudy.org), held in the NIMH Data Archive (NDA). This is a multisite, longitudinal study designed to recruit more than 10,000 children age 9-10 and follow them over 10 years into early adulthood. The ABCD Study® is supported by the National Institutes of Health and additional federal partners under award numbers U01DA041048, U01DA050989, U01DA051016, U01DA041022, U01DA051018, U01DA051037, U01DA050987, U01DA041174, U01DA041106, U01DA041117, U01DA041028, U01DA041134, U01DA050988, U01DA051039, U01DA041156, U01DA041025, U01DA041120, U01DA051038, U01DA041148, U01DA041093, U01DA041089, U24DA041123, U24DA041147. A full list of supporters is available at https://abcdstudy.org/federal-partners.html. A listing of participating sites and a complete listing of the study investigators can be found at https://abcdstudy.org/consortium_members/. ABCD consortium investigators designed and implemented the study and/or provided data but did not necessarily participate in the analysis or writing of this report. This manuscript reflects the views of the authors and may not reflect the opinions or views of the NIH or ABCD consortium investigators.

## Supplementary Materials: Figure S1

**Figure S1:**
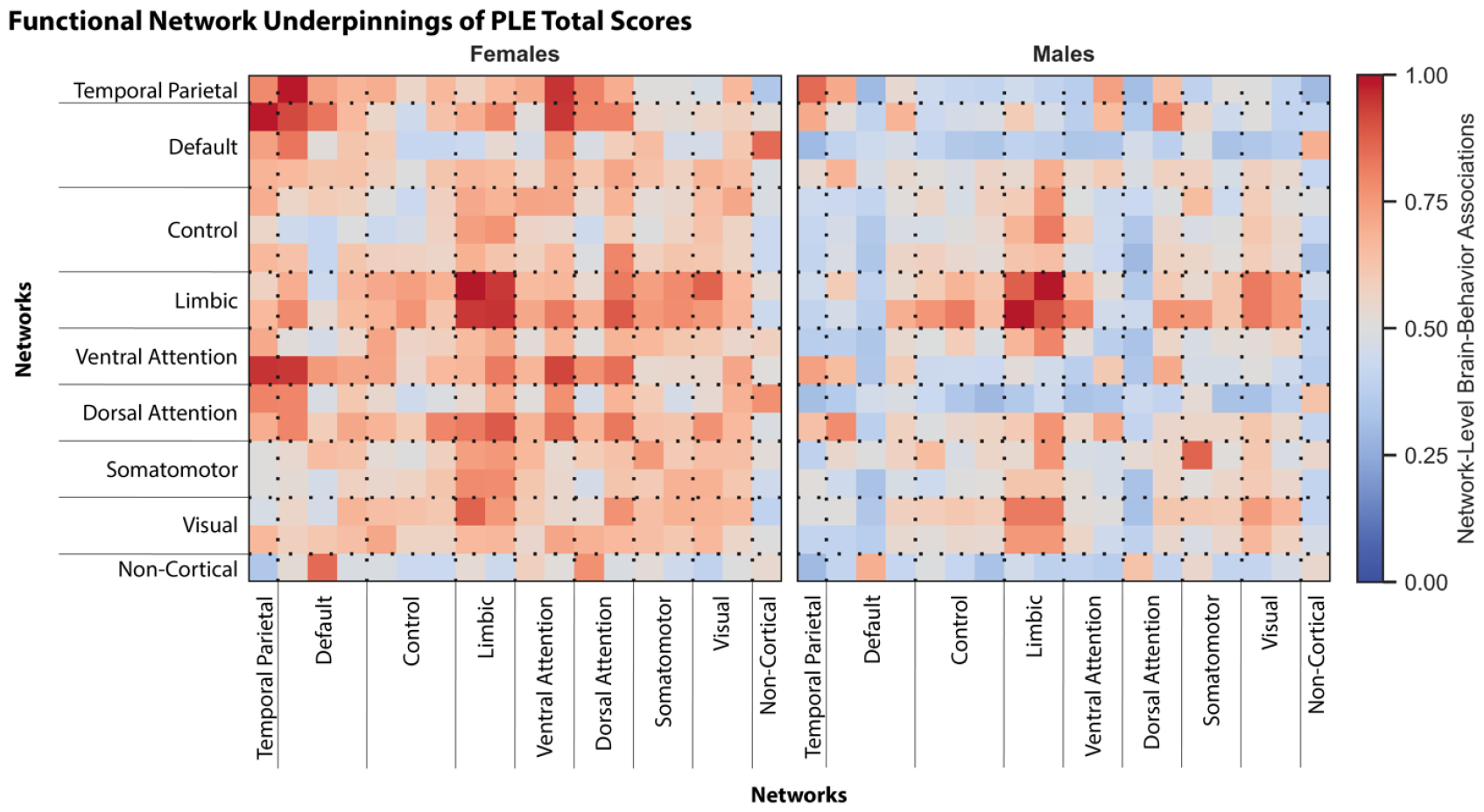
Overlapping and distinct functional networks underlie PLEs across the sexes. Absolute associations between functional network connectivity and PLE Total scores in assigned females (left) and males (right) are shown as per the colormap. Warmer colors indicate stronger correlations and cooler colors indicate weaker correlations. To facilitate visualization, values within each matrix were divided by the maximum value within that matrix.

## References

1. Lieberman, J.A. and M.B. First, Psychotic disorders. New England Journal of Medicine, 2018. 379(3): p. 270–280.

2. Karcher, N.R., Psychotic-like experiences in childhood and early adolescence: Clarifying the construct and future directions. Schizophrenia research, 2022. 246: p. 205–206.

3. Karcher, N.R., et al., Assessment of the Prodromal Questionnaire–Brief Child Version for measurement of self-reported psychoticlike experiences in childhood. JAMA psychiatry, 2018. 75(8): p. 853–861.

4. Barnes, G., et al., Distressing psychotic-like experiences, cognitive functioning and early developmental markers in clinically referred young people aged 8–18 years. Social Psychiatry and Psychiatric Epidemiology, 2022: p. 1–12.

5. Fisher, H., et al., Specificity of childhood psychotic symptoms for predicting schizophrenia by 38 years of age: a birth cohort study. Psychological medicine, 2013. 43(10): p. 2077–2086.

6. Riecher-Rössler, A., Sex and gender differences in mental disorders. The Lancet Psychiatry, 2017. 4(1): p. 8–9.

7. Riecher-Rössler, A., S. Butler, and J. Kulkarni, Sex and gender differences in schizophrenic psychoses—a critical review. Archives of women’s mental health, 2018. 21(6): p. 627–648.

8. Eaton, N.R., et al., An invariant dimensional liability model of gender differences in mental disorder prevalence: evidence from a national sample. Journal of abnormal psychology, 2012. 121(1): p. 282.

9. Li, R., et al., Why sex differences in schizophrenia? Journal of translational neuroscience, 2016. 1(1): p. 37.

10. Wu, Z., et al., Sex difference in the prevalence of psychotic-like experiences in adolescents: results from a pooled study of 21,248 Chinese participants. Psychiatry Research, 2022. 317: p. 114894.

11. Sha, Z., et al., Common dysfunction of large-scale neurocognitive networks across psychiatric disorders. Biological psychiatry, 2019. 85(5): p. 379–388.

12. Cole, M.W., G. Repovš, and A. Anticevic, The frontoparietal control system: a central role in mental health. The Neuroscientist, 2014. 20(6): p. 652–664.

13. Chen, J., et al., Shared and unique brain network features predict cognitive, personality, and mental health scores in the ABCD study. Nat Commun, 2022. 13(1): p. 2217.

14. Shanmugan, S., et al., Sex differences in the functional topography of association networks in youth. Proceedings of the National Academy of Sciences, 2022. 119(33): p. e2110416119.

15. Dhamala, E., et al., Functional brain networks are associated with both sex and gender in children. bioRxiv, 2023.

16. Dhamala, E., et al., Brain-based predictions of psychiatric illness-linked behaviors across the sexes. Biological Psychiatry, 2023.

17. Loewy, R.L., et al., Psychosis risk screening with the Prodromal Questionnaire—brief version (PQ-B). Schizophrenia research, 2011. 129(1): p. 42–46.

18. Cicero, D.C., A. Krieg, and E.A. Martin, Measurement invariance of the Prodromal Questionnaire–Brief among White, Asian, Hispanic, and multiracial populations. Assessment, 2019. 26(2): p. 294–304.

19. Karcher, N.R., et al., Replication of associations with psychotic-like experiences in middle childhood from the adolescent brain cognitive development (ABCD) study. Schizophrenia Bulletin Open, 2020. 1(1): p. sgaa009.

20. Chen, J., et al., Relationship between prediction accuracy and feature importance reliability: An empirical and theoretical study. NeuroImage, 2023. 274: p. 120115.

21. Li, J., et al., Global signal regression strengthens association between resting-state functional connectivity and behavior. Neuroimage, 2019. 196: p. 126–141.

22. Haufe, S., et al., On the interpretation of weight vectors of linear models in multivariate neuroimaging. Neuroimage, 2014. 87: p. 96–110.

23. Yeo, B.T., et al., The organization of the human cerebral cortex estimated by intrinsic functional connectivity. J Neurophysiol, 2011. 106(3): p. 1125–65.

24. Karcher, N.R., et al., Resting-state functional connectivity and psychotic-like experiences in childhood: results from the adolescent brain cognitive development study. Biological psychiatry, 2019. 86(1): p. 7–15.

25. Kelleher, I., et al., Clinicopathological significance of psychotic experiences in non-psychotic young people: evidence from four population-based studies. The British Journal of Psychiatry, 2012. 201(1): p. 26–32.

26. Maitra, R., et al., Psychotic like experiences in healthy adolescents are underpinned by lower fronto-temporal cortical gyrification: a study from the IMAGEN consortium. Schizophrenia Bulletin, 2023. 49(2): p. 309–318.

27. Karcher, N.R., et al., Persistent and distressing psychotic-like experiences using adolescent brain cognitive developmentlZI study data. Molecular Psychiatry, 2022. 27(3): p. 1490–1501.

28. Shapiro, J.R., S.L. Klein, and R. Morgan, Stop ‘controlling’for sex and gender in global health research. BMJ Global Health, 2021. 6(4): p. e005714.

29. Meyer, G.J., et al., Psychological testing and psychological assessment: A review of evidence and issues. American psychologist, 2001. 56(2): p. 128.

30. Kaymaz, N., et al., Do subthreshold psychotic experiences predict clinical outcomes in unselected non-help-seeking population-based samples? A systematic review and meta-analysis, enriched with new results. Psychological medicine, 2012. 42(11): p. 2239–2253.

31. Healy, C., et al., Childhood and adolescent psychotic experiences and risk of mental disorder: a systematic review and meta-analysis. Psychological medicine, 2019. 49(10): p. 1589–1599.

32. Maddox, L., et al., Cognitive behavioural therapy for unusual experiences in children: a case series. Behavioural and cognitive psychotherapy, 2013. 41(3): p. 344–358.

33. Addington, J., et al., North American Prodrome Longitudinal Study: a collaborative multisite approach to prodromal schizophrenia research. Schizophrenia bulletin, 2007. 33(3): p. 665–672.

34. Addington, J., et al., North American prodrome longitudinal study (NAPLS 2): overview and recruitment. Schizophrenia research, 2012. 142(1-3): p. 77–82.

35. Addington, J., et al., North American prodrome longitudinal study (NAPLS 3): methods and baseline description. Schizophrenia research, 2022. 243: p. 262–267.

36. Li, X., W. Zhou, and Z. Yi, A glimpse of gender differences in schizophrenia. General Psychiatry, 2022. 35(4).

37. Culbert, K.M., K.N. Thakkar, and K.L. Klump, Risk for midlife psychosis in women: critical gaps and opportunities in exploring perimenopause and ovarian hormones as mechanisms of risk. Psychological medicine, 2022. 52(9): p. 1612–1620.

38. Cohen, R.Z., et al., Earlier puberty as a predictor of later onset of schizophrenia in women. American Journal of Psychiatry, 1999. 156(7): p. 1059–1065.

39. Sydnor, V.J., et al., Neurodevelopment of the association cortices: Patterns, mechanisms, and implications for psychopathology. Neuron, 2021. 109(18): p. 2820–2846.

40. De Bellis, M.D., et al., Sex differences in brain maturation during childhood and adolescence. Cereb Cortex, 2001. 11(6): p. 552–7.

41. Gur, R.E. and R.C. Gur, Sex differences in brain and behavior in adolescence: Findings from the Philadelphia Neurodevelopmental Cohort. Neurosci Biobehav Rev, 2016. 70: p. 159–170.

42. Ricard, J., et al., Confronting racially exclusionary practices in the acquisition and analyses of neuroimaging data. Nature Neuroscience, 2022: p. 1–8.

43. Wierenga, L.M., et al., Recommendations for a better understanding of sex and gender in neuroscience of mental health. Biological Psychiatry Global Open Science, 2023: p. 100283.

44. Dhamala, E., et al., Distinct functional and structural connections predict crystallised and fluid cognition in healthy adults. Hum Brain Mapp, 2021. 42(10): p. 3102–3118.

45. Dhamala, E., et al., Shared functional connections within and between cortical networks predict cognitive abilities in adult males and females. Hum Brain Mapp, 2022.

46. Dhamala, E., et al., Proportional intracranial volume correction differentially biases behavioral predictions across neuroanatomical features and populations. NeuroImage, 2022.

47. Casey, B.J., et al., The Adolescent Brain Cognitive Development (ABCD) study: Imaging acquisition across 21 sites. Dev Cogn Neurosci, 2018. 32: p. 43–54.

48. Ooi, L.Q.R., et al., Comparison of individualized behavioral predictions across anatomical, diffusion and functional connectivity MRI. BioRxiv, 2022.

49. Hagler, D.J., Jr., et al., Image processing and analysis methods for the Adolescent Brain Cognitive Development Study. Neuroimage, 2019. 202: p. 116091.

50. Gordon, E.M., et al., Generation and evaluation of a cortical area parcellation from resting-state correlations. Cerebral cortex, 2016. 26(1): p. 288–303.

51. Kong, R., et al., Spatial Topography of Individual-Specific Cortical Networks Predicts Human Cognition, Personality, and Emotion. Cerebral Cortex, 2018. 29(6): p. 2533–2551.

52. Potter, A.S., et al., Measurement of gender and sexuality in the Adolescent Brain Cognitive Development (ABCD) study. Developmental Cognitive Neuroscience, 2022. 53: p. 101057.

53. Benjamini, Y. and Y. Hochberg, Controlling the False Discovery Rate - a Practical and Powerful Approach to Multiple Testing. Journal of the Royal Statistical Society Series B-Statistical Methodology, 1995. 57(1): p. 289–300.

54. Chen, J., et al., There is no fundamental trade-off between prediction accuracy and feature importance reliability. bioRxiv, 2022.

55. Dale, A.M., B. Fischl, and M.I. Sereno, Cortical surface-based analysis. I. Segmentation and surface reconstruction. Neuroimage, 1999. 9(2): p. 179–94.

56. Fischl, B., A. Liu, and A.M. Dale, Automated manifold surgery: constructing geometrically accurate and topologically correct models of the human cerebral cortex. IEEE Trans Med Imaging, 2001. 20(1): p. 70–80.

57. Fischl, B., M.I. Sereno, and A.M. Dale, Cortical surface-based analysis. II: Inflation, flattening, and a surface-based coordinate system. Neuroimage, 1999. 9(2): p. 195–207.

58. Fischl, B., et al., High-resolution intersubject averaging and a coordinate system for the cortical surface. Hum Brain Mapp, 1999. 8(4): p. 272–84.

59. Greve, D.N. and B. Fischl, Accurate and robust brain image alignment using boundary-based registration. Neuroimage, 2009. 48(1): p. 63–72.

60. Jenkinson, M., et al., Improved optimization for the robust and accurate linear registration and motion correction of brain images. Neuroimage, 2002. 17(2): p. 825–841.

61. Power, J.D., et al., Spurious but systematic correlations in functional connectivity MRI networks arise from subject motion. Neuroimage, 2012. 59(3): p. 2142–54.

62. Power, J.D., et al., Distinctions among real and apparent respiratory motions in human fMRI data. NeuroImage, 2019. 201: p. 116041.

63. Fair, D.A., et al., Correction of respiratory artifacts in MRI head motion estimates. Neuroimage, 2020. 208: p. 116400.

64. Gratton, C., et al., Removal of high frequency contamination from motion estimates in single-band fMRI saves data without biasing functional connectivity. Neuroimage, 2020. 217: p. 116866.

65. Greene, A.S., et al., Task-induced brain state manipulation improves prediction of individual traits. Nature communications, 2018. 9(1): p. 1–13.

66. Li, J.W., et al., Global signal regression strengthens association between resting-state functional connectivity and behavior. Neuroimage, 2019. 196: p. 126–141.

67. Power, J.D., et al., Methods to detect, characterize, and remove motion artifact in resting state fMRI. Neuroimage, 2014. 84: p. 320–41.

68. Schaefer, A., et al., Local-global parcellation of the human cerebral cortex from intrinsic functional connectivity MRI. Cerebral cortex, 2018. 28(9): p. 3095–3114.

69. Fischl, B., et al., Whole brain segmentation: automated labeling of neuroanatomical structures in the human brain. Neuron, 2002. 33(3): p. 341–55.

70. He, T., et al., Deep neural networks and kernel regression achieve comparable accuracies for functional connectivity prediction of behavior and demographics. NeuroImage, 2020. 206: p. 116276.

71. Dhamala, E., B.T. Yeo, and A.J. Holmes, Methodological Considerations for Brain-Based Predictive Modelling in Psychiatry. Biological Psychiatry, 2022.

72. Tian, Y. and A. Zalesky, Machine learning prediction of cognition from functional connectivity: Are feature weights reliable? bioRxiv, 2021.

73. Yeo, B.T., et al., The organization of the human cerebral cortex estimated by intrinsic functional connectivity. Journal of neurophysiology, 2011. 106(3): p. 1125–1165.

